# Drug-gene interaction screens coupled to tumour data analyses identify the most clinically-relevant cancer vulnerabilities driving sensitivity to PARP inhibition

**DOI:** 10.1101/2022.07.29.501846

**Authors:** Kunzah Jamal, Alessandro Galbiati, Joshua Armenia, Giuditta Illuzzi, James Hall, Sabrina Bentouati, Daniel Barrell, Miika Ahdesmäki, Functional Genomics Centre, Mark J. O’Connor, Elisabetta Leo, Josep V. Forment

**Author notes:** Corresponding author: Dr Josep Forment, DDR Biology, Bioscience, Oncology R&D, AstraZeneca, Chesterford Research Park, Cambridge CB10 1XL, UK. These authors contributed equally to this work (Kunzah Jamal, Alessandro Galbiati, Joshua Armenia). Artios Pharma Limited, B940, Babraham Research Campus, Cambridge, CB22 3FH, UK.

## Abstract

Poly(ADP-ribose) polymerase (PARP) inhibitors (PARPi) are currently indicated for the treatment of ovarian, breast, pancreatic and prostate cancers harbouring mutations in the tumour suppressor genes *BRCA1* or *BRCA2*. In the case of ovarian and prostate cancers, their classification as homologous recombination repair (HRR) deficient (HRD) or mutated (HRRm) also makes PARPi an available treatment option beyond *BRCA1* or *BRCA2* mutational status. However, identification of the most relevant genetic alterations driving the HRD phenotype has proven difficult and recent data have shown that other genetic alterations not affecting HRR are also capable of driving PARPi responses. To gain insight into the genetics driving PARPi sensitivity, we performed CRISPR-Cas9 loss-of-function screens in 6 PARPi-insensitive cell lines and combined the output with published PARPi datasets from 8 additional cell lines. Ensuing exploration of the data identified 110 genes whose inactivation is strongly linked to sensitivity to PARPi. Parallel cell line generation of isogenic gene knockouts in ovarian and prostate cancer cell lines identified that inactivation of core HRR factors is required for driving *in vitro* PARPi responses comparable to the ones observed for *BRCA1* or *BRCA2* mutations. Moreover, pan-cancer genetic, transcriptomic and epigenetic data analyses of these 110 genes highlight the ones most frequently inactivated in tumours, making this study a valuable resource for prospective identification of potential PARPi-responsive patient populations. Importantly, our investigations uncover *XRCC3* gene silencing as a potential new prognostic biomarker of PARPi sensitivity in prostate cancer.

**Statement of significance:** This study identifies tumour genetic backgrounds where to expand the use of PARP inhibitors beyond mutations in *BRCA1* or *BRCA2*. This is achieved by combining the output of unbiased genome-wide loss-of-function CRISPR-Cas9 genetic screens with bioinformatics analysis of biallelic losses of the identified genes in public tumour datasets, unveiling loss of the DNA repair gene *XRCC3* as a potential biomarker of PARP inhibitor sensitivity in prostate cancer.

## INTRODUCTION

The breast cancer susceptibility genes *BRCA1* and *BRCA2* (*BRCA* genes) are well-known tumour suppressors whose inactivation increases the probability of developing cancer, particularly of breast and ovarian origin (1), but also in the pancreas and prostate (2,3). BRCA proteins are key components of the homologous recombination repair (HRR) pathway of DNA repair and their deficiency fosters genome instability, one of the hallmarks of cancer (reviewed in (4)). Poly(ADP-ribose) polymerase (PARP) inhibitors (PARPi) are current treatment options for ovarian, breast, pancreatic and prostate cancer patients harbouring mutations in *BRCA1* or *BRCA2* (BRCAm) but they have also been approved in broader patient populations in ovarian and prostate cancer (5-7). These broader approvals have involved the identification of tumours with HRR deficiency (HRD) beyond BRCAm, either by the use of genomic detection of genome instability patterns (“genomic scars”) linked to HRD or by genetic identification of mutations in non-BRCA genes linked to HRR (HRRm) (8,9).

Different HRRm gene panels have been used in clinical trials to identify the best biomarkers of response to PARPi. Most of these efforts have almost invariably found that HRRm beyond *BRCA* genes are rare, making it difficult to assess the validity of some of these biomarkers (7,10,11). However, there seems to be a significant HRD tumour population identified through detection of genomic scars for which current HRRm gene panels fail to explain their genetic origin (5,6). Moreover, recent pre-clinical work has highlighted that mutations in genes not involved in HRR could be valid biomarkers to predict PARPi sensitivity (12-14), probably suggesting the need of a more holistic gene selection strategy in these panels.

In this work, and in an effort to identify cancer-relevant genetic biomarkers of PARPi responses in an unbiased way, we carried out CRISPR-Cas9 genome-wide loss-of-function (LoF) screens (reviewed in (15)) to uncover genetic determinants of sensitivity to PARPi in a variety of cell lines. We coupled the output with a bespoke analysis pipeline of tumour genetic and epigenetic data to uncover the most clinically-relevant vulnerabilities to PARPi, identifying potential new biomarkers of PARPi sensitivity. In addition, we provide for the first time head-to-head comparison of PARPi responses between defects in *BRCA* genes and other genes identified by our analyses through generation of isogenic cell line pairs in clinically-relevant tissue types, namely ovarian and prostate cancer cells.

## MATERIALS AND METHODS

### Cell lines and compounds

LNCAP, DU145 (NCI-DTP Cat# DU-145, RRID:CVCL_0105) and SKOV3 (NCI-DTP Cat# SKOV-3, RRID:CVCL_0532) cells were obtained from ATCC. The DLD1 wild type and *BRCA2* -/-cell lines were purchased from Horizon Discovery. Cell line identification (STR typing) was validated using the CellCheck assay (IDEXX Bioanalytics, Westbrook, ME, USA). All cell lines were validated free of virus mycoplasma contamination using the MycoSEQ assay (Thermo Fisher Scientific, Waltham, MA, USA) or STAT-Myco assay (IDEXX Bioanalytics). All cell lines were grown according to supplier instructions. LNCAP, DU145 and DLD1 were grown in RPMI-1640 growth media (Corning 17-105-CV) supplemented with 10% fetal bovine serum (FBS) and 2 mM glutamine. SKOV3 were grown in McCoy’s 5A (Modified) Medium (ThermoFisher Scientific 16600082). Olaparib and AZD0156 (ATM inhibitor, ATMi) were made by AstraZeneca (Cambridge, UK), carboplatin and cisplatin were bought from Tocris Bioscience (cat. No. 2626 and 15663-27-1). Olaparib, ATMi and carboplatin were all solubilised in DMSO at 10 mM stock concentration. Cisplatin was solubilised in an aqueous solution at 1.67___mM.

### Gene expression analysis by RT-qPCR

Total RNA was isolated from cells in 96-well plates using the Qiagen FastLane Cell Probe Kit (QIAGEN, 216413), according to the manufacturer’s instructions to a final volume of 40 μL per well, at the indicated time point after treatment. Gene expression was evaluated by qPCR using the ONE-step QuantiTect Probe RT-PCR Kit (Qiagen, Cat No./ID: 204445). For each reaction 2 μL of RNA were used. Real-time qPCR reactions were performed on a Roche Lightcycler 480 II Sequence Detection System.

The following Taqman probes were obtained from ThermoFisher Scientific: BRCA2 (Hs00609073_m1), IPO8 (Hs00914057_m1), RAD51B (Hs01568768_m1), RAD54L (Hs00941668.m1), Actin (Hs01060665.g1).

For data analysis dCt was calculated by subtracting Average Ct of housekeeping genes from each Ct. An average dCt for the control group was calculated and subtracted from dCt to calculate negative ddCt. 2^negative ddCt was used to calculate Fold Change.

### Immunoblotting

Cells were lysed in RIPA buffer (Sigma-Aldrich) supplemented with protease inhibitors (Roche, Basel, Switzerland), phosphatase inhibitors (Sigma-Aldrich) and benzonase (Merck, cat. No. 103773). After 30 minutes (min) of incubation on ice the lysates were cleared through centrifugation at 15,000 rpm 4°C for 20 min and supernatant were kept for sample loading. NuPAGE™LDS Sample Buffer (ThermoFisher Scientific, cat. No. NP0008) and NuPAGE™Sample Reducing Agent (ThermoFisher Scientific, cat. No. NP0004) were added to the samples. Equal amounts of whole cell lysates were separated on 4-12% Bis-Tris NuPAGE gels and analysed by standard immunoblotting. Immunoblots are representative of experiments that were performed at least twice. For ATM signalling pathway analysis, cells were pre-treated with the ATMi for 1hr prior to ionizing radiation induction by a high-voltage X-ray-generator tube (Faxitron X-Ray Corporation).

The following antibodies were used:

| Protein | Species | Cat number | Supplier | Dilution |
| --- | --- | --- | --- | --- |
| Actin | mouse | A2228 | Sigma | 1:2000 |
| ATM | mouse | ab78 | Abcam | 1:2000 |
| pATM (S1981) | rabbit | ab81282 | Abcam | 1:1000 |
| BRCA1 | mouse | OP92 | Merck Millipore | 1:1000 |
| Cas9 | rabbit | ab189380 | Abcam | 1:1000 |
| GAPDH | Rabbit | 2118 | Cell Signalling | 1:1000 |
| KAP1 | rabbit | ab10483 | Abcam | 1:1000 |
| pKAP1 (S824) | rabbit | ab70483 | Abcam | 1:1000 |
| pKAP1 (S473) | rabbit | 654102 | BioLegend | 1:1000 |
| pCHEK2 | rabbit | 2661 | Cell Signalling | 1:1000 |
| KAP1 | rabbit | ab10483 | Abcam | 1:1000 |
| pKAP1 (S824) | rabbit | ab70483 | Abcam | 1:1000 |
| pKAP1 (S473) | rabbit | 654102 | BioLegend | 1:1000 |
| PALB2 | rabbit | A301-246A | Bethyl Laboratories | 1:1000 |
| RAD51C | mouse | NB100-177 | Novus | 1:1000 |

### Immunofluorescence – RAD51 foci assay

Cells were plated in 96-well plates (Perkin Elmer CellCarrier-96 Ultra Microplates, cat. No. 6055302) to reach 70% confluency the next day and ionizing radiation was induced by a high-voltage X-ray-generator tube (Faxitron X-Ray Corporation). For EdU staining, cells were incubated with 10 µM EdU for 1 h prior to fixation. At the indicated time points after treatment, cells were fixed in 4% PFA for 15 min at RT and then permeabilized in PBS+0.1% Triton X-100 for 10 min at RT. Blocking was performed using 0.5% BSA + 0.2% gelatin from cold water fish skin (Sigma, cat no. G7765) in PBS for 1 h at RT. Cells were washed with PBS and incubated with EdU click-It reaction (HEPES pH 7.5 125mM, CuSO4·5H2O 20mM, Ascorbate 100mM, Alexa Azide 647 (Sigma, A10277) 5mM), for 20 min at RT in the dark. The Click-it reaction was removed and cells were washed 3 times with PBG. Primary antibodies were incubated overnight at 4°C, followed by Alexa-Fluor secondary antibodies and DAPI (Sigma, 1 µg/ml) for 1 h at RT. The following antibodies were used:

|  | Species | Cat number | Supplier | Dilution |
| --- | --- | --- | --- | --- |
| Alexa Fluor 488 secondary | rabbit | A11037 | Thermo | 1:2000 |
| Alexa Fluor 594 secondary | mouse | A21099 | Thermo | 1:2000 |
| γH2AX | mouse | 05-636 | Merck Millipore | 1:2000 |
| RAD51 | rabbit | 70-001 | Bioacademia | 1:7000 |

### Colony formation assay (CFA)

To evaluate the efficacy of olaparib and platinum, a colony formation assay was employed. Here, cells were seeded at low density in a 24-well plate and exposed to olaparib for a time corresponding to more than 5 replication cell cycles: all except BRCA2, PALB2 and RAD51C KO cells were plated at 500 cells per well for 9-14 days whereas BRCA2, PALB2 and RAD51C KO cells were plated at 800 cells per well for 10-14 days. After cell attachment, compounds were dispensed with automated digital D300 HP dispenser (Tecan). Drugs were added from compound stocks dissolved in DMSO, in 7 titration dilutions, where each concentration was tested in triplicate in each plate. DMSO served as a vehicle control. The concentration ranges tested were chosen to obtain dose-response and to cover minimal to maximal activity of a given compound. Plates were incubated at 37°C, 5% CO2 for indicated times. Colony formation and absence of contamination was regularly checked on the microscope. Next, colonies were fixed and stained using Blue-G-250 brilliant blue (#B8522-1EA, Sigma, reconstituted in 25% (v/v) methanol and 5% (v/v) acetic acid) for 15 minutes. Prior to imaging, plates were thoroughly washed with de-ionised water (dH2O). Plates with stained colonies were scanned with GelCount™(Oxford OPTRONIX) at 600 dpi resolution. Colony formation was scored by quantifying the total optical density measured with ImageJ software (RRID:SCR_003070), using a 24-well plate Region of Interest (ROI) mask. Data analysis was performed by normalisation on the vehicle treated of the respective plate set as = 100. Data were normalised and plotted to respective vehicle-control (set as = 100) and IC50 were calculated using Prism GraphPad software.

### Cell proliferation assay – CellTiter-Glo

To evaluate the efficacy of olaparib and platinum, a CellTiter-glo assay was employed. Here, cells were seeded at low density (400-500 cells per well) in a 96-well plate and exposed to olaparib for 7-8 days allowing the cells to go through at least 5 replication cycles. After cell attachment, compounds were dispensed with automated digital D300 HP dispenser (Tecan). Drugs were added from compound stocks dissolved in DMSO, in 7 titration dilutions, where each concentration was tested in a triplicate in each plate. DMSO served as a vehicle control. The concentration ranges tested were chosen to obtain dose-response and to cover minimal to maximal activity of a given compound. Plates were incubated at 37ºC, 5% CO2 for indicated times. Cell growth was stopped by adding CellTiter-Glo as per manufacturer’s instructions (Promega, Madison, WI, USA; G7570). Data analysis was performed by normalisation on the vehicle treated of the respective plate set as = 100. Data were normalised and plotted to respective vehicle-control (set as = 100) using Prism GraphPad software.

### Lentiviral production and transduction

HEK293T cells seeded at the concentration of 400,000 cells per well in 6-well plates were transfected with the Cas9 expressing plasmid or the sgRNA targeting plasmid and viral packaging, psPax2 and pMD2.G at the following mixing ratio: 0.9 _μ_g lentiviral vector (Cas9 or sgRNA cloned into vector), 0.9 _μ_g psPax2 and 0.2 _μ_g pMD2.G, 2uL PLUS reagent, 6 _μ_L Lipofectamine LTX (ThermoFisher Scientific, cat. No. 15338100) in 500 _μ_L per well. The reaction mix was incubated for 30 min at RT and then added to the cells in 1.5 mL Opti-MEM. 6 h after transfection, OPTIMEM was replaced with fresh medium and 72 h after transfection the virus-containing supernatant was collected and filtered with 0.45 _μ_M filters.

### Generation of Cas9 expressing cells

To produce SKOV3 cells stably expressing Cas9, these were transduced with the pKLV2-EF1a-BsdCas9-W expressing vector. To establish the LNCAP cell line expressing a doxycycline-inducible Cas9, cells were transduced with the pBSK-TOIC-Cas9-T2A-TagBFP-Obl-r26-AAVS-Invneo vector. Following transduction, cells were selected with Blasticidine at 15 µg/mL for 3 passages and analysed by western blot.

### sgRNA cloning into lentiviral vector

For the SKOV3, DLD1 and LNCAP KO generation, the sgRNA targeting the sequence of interest was cloned into the Bbsl region of the vector pKLV-U6gRNA(BbsI)-PGKpuro2ABFP or pKLV2-hU6gRNA5(BbsI)-EF1a-mClover3-T2A-HygR-W, as previously described in (16). Correct cloning was verified via Sanger sequencing.

The following sgRNA sequences were used for cell line generation:

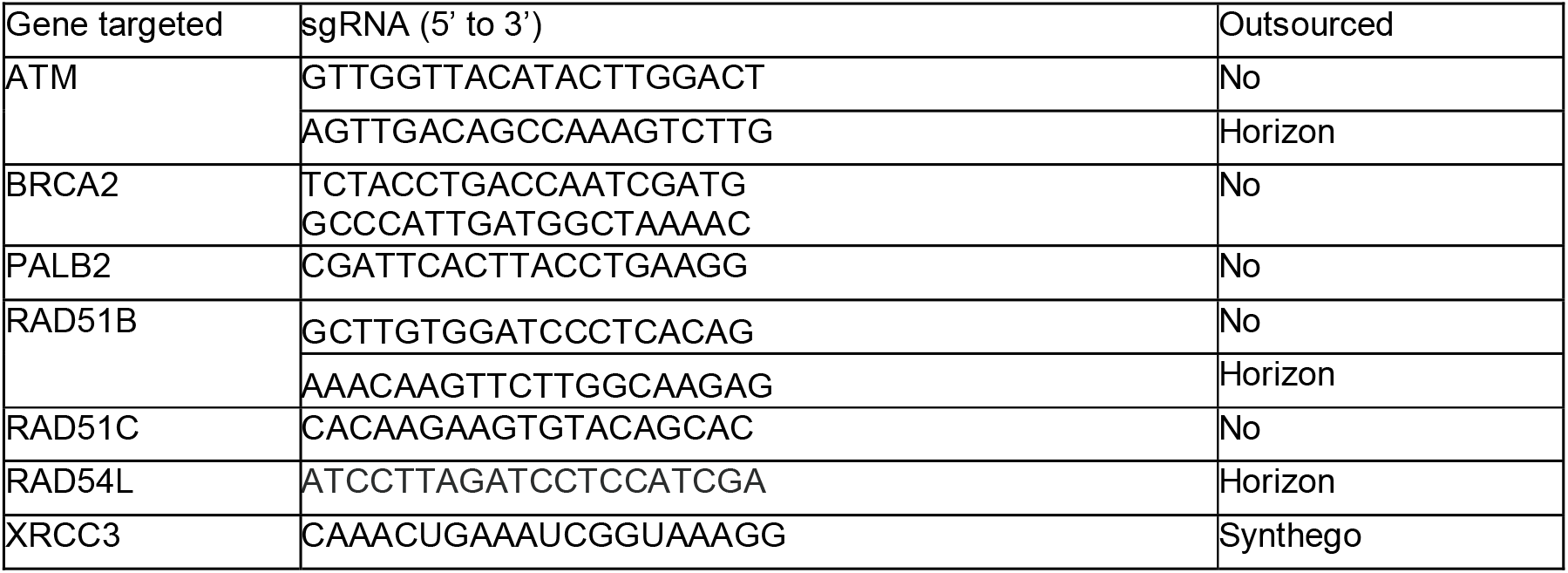

### Generation of CRISPR knockout cell lines

To generate the CRISPR/Cas9 KO cell lines, cells were seeded at 100,000 cells per well in a 6 well plate. For the LNCAP cells, Cas9 was induced with 100 ng/mL doxycyclin (Sigma) to allow Cas9 expression and after 24 hrs the cell medium was refreshed without doxycyclin. The next day, cells were transduced with the lentivirus containing the sgRNA targeting the gene of interest or a non-targeting sgRNA (control, CTRL, cell line) as described above. Viral supernatant was mixed in 2mL cell culture medium supplemented with 8 _μ_g/ml polybrene (Millipore) and further incubated overnight at 37°C. The medium was refreshed on the following day and the transduced cells were cultured further. 48 h following transduction, transduced cells were moved into antibiotic selection (Hygromycin, 100 µg/ml or puromycin, 0.5 µg/ml) for 5 days. Cell lines were expanded and the KO of the gene of interest was validated by immunoblot, TIDE or ICE sequencing (17,18) or an RT-PCR to assess the KO efficiency. Single cell clones were derived from the KO pools from serial dilution plating in 96-well plates and KO efficiency was validated with western blot, TIDE or RT-PCR.

### Statistical analyses

Results are shown as mean ± standard error of the mean (s.e.m) or percentages ± 95% confidence interval (c.i.) as indicated. P-value was calculated by Student’s two-tailed t-test or chi-squared test, respectively, using Prism software. Mutual exclusivity analysis was performed using Fisher’s exact test.

### CRISPR screens and data analysis

CAMA1, U2OS, DLD1, T47D, HCC1806 (KCB Cat# KCB 2014032YJ, RRID:CVCL_1258) and BT549 (NCI-DTP Cat# BT-549, RRID:CVCL_1092) breast cancer cells were infected with lentiviral particles containing the whole-genome sgRNA library, subjected to puromycin selection, and passaged to ensure loss of affected protein products. Puromycin-resistant cells were exposed to 1 μM olaparib for 21 days, and resistant pools were isolated. Genomic DNA was extracted from these and from parallel cell cultures treated in the absence of olaparib, and DNA libraries were prepared and sequenced. Genomic DNA was extracted and gRNAs sequenced as described previously (16). Single-end Illumina sequencing reads of 19 nucleotides were counted for each gRNA using in-house written software.

For all the screens each replicate of plasmid, DMSO control and PARPi-treated (or *PARP1* KO) cell line sequenced samples were first counted for exact matches of the Yusa library (19). The counts were then analysed for negative selection i.e. depletion in the treatment vs control replicates (2 each) by the MAGeCK algorithm (20). Genes whose false discovery rate was less than or equal to 0.1 were included as sensitising hits. In addition, each DMSO control sample was further assessed against the plasmid sample using BAGEL (21) to ensure the expected depletion of essential genes as a quality control measure.

### Biallelic inactivation analysis in TCGA data

Biallelic inactivation prevalence was estimated from TCGA pancancer data by evaluating prevalence of somatic/germline alterations using internal variant calls (22,23), homozygous deletion and promoter methylation data using level 3 data from TCGA. Biallelic inactivation was defined as (i) a germline pathogenic mutation with loss of heterozygosity (LOH) of the wild-type allele, (ii) a germline pathogenic mutation and a somatic pathogenic mutation, (iii) a somatic pathogenic mutation with LOH of the wild-type allele, (iv) two different somatic pathogenic mutations, (v) homozygous deletion or (vi) promoter hypermethylation.

### Data availability statement

The data generated in this study are available within the article and its supplementary data files. Raw data from the CRISPR screens performed in this study can be found at the European Nucleotide Archive (ENA) at EMBL-EBI under accession number PRJEB54620.

## RESULTS

### CRISPR screens to identify genes involved in the response to PARPi

As a way to uncover potential biomarkers of response to PARPi in an unbiased way, we performed genome-wide CRISPR-Cas9 loss-of-function (LoF) screens in 6 tumour cell lines from different tissue of origin (breast, colon and bone), looking for genes whose inactivation would render cells sensitive to PARPi. The 6 cell lines were chosen by them carrying no genetic or epigenetic alterations in *BRCA* or other HRR genes and presenting a half-maximal inhibitory concentration (IC50) for olaparib at least 10-fold higher than the values reported for *BRCAm* cell lines in colony-forming assays (in the 10-100 nM range) (24) to maximize the chances of identifying sensitization events. The output was combined with results from published CRISPR screens in 8 other cell lines from different tissue of origin (cervix, retina, breast, ovary, skin, blood, kidney and colon) that, except one (SUM149PT, a *BRCA1m* breastcancer cell line), do not carry genetic or epigenetic alterations in *BRCA* or other HRR genes (12,25-28) (**Fig 1A**). To standardize the results, all raw data (where available) were run through the same analysis pipeline (see Methods), which identified 1147 genes whose loss would confer sensitivity to PARPi (or compromised fitness in a PARP1 deficient background; **Supp Table S1**). As a way to generate a list of high confidence genes, we selected genes that were identified in more than one independent screen, which reduced the number of genes in the list to 110 (**Fig 1B; Supp Table S2**). Pathway analysis of these 110 genes revealed a strong enrichment in DNA repair genes, especially in genes involved in HRR, as expected by the ranking of genes that were identified more often in all screens analysed (**Fig 1C-D**).

**Figure 1.**
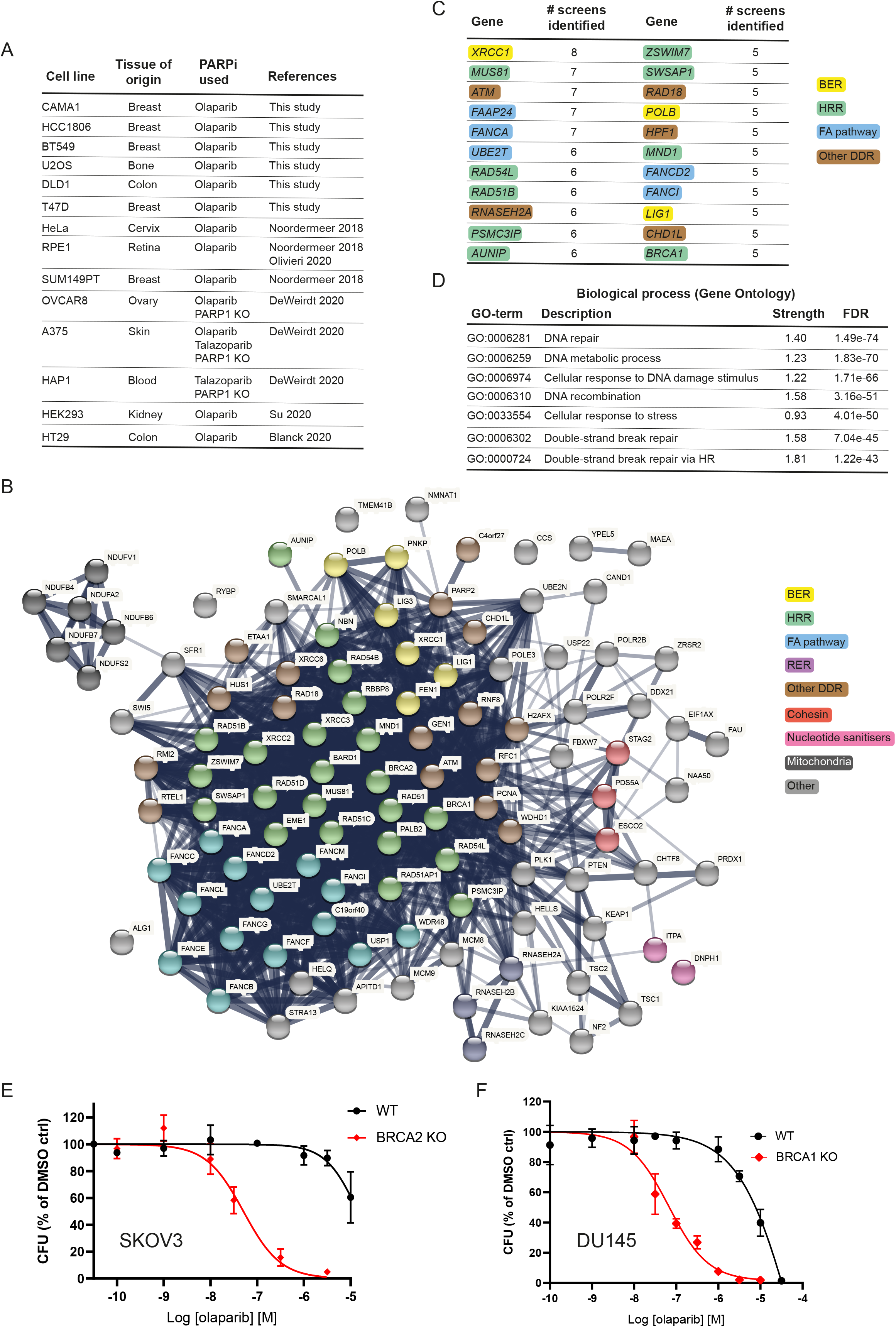
Identification of biomarkers of PARPi sensitivity through CRISPR-Cas9 loss-of-function screens. **A**, Summary of cell lines and CRISPR screens performed or analysed in this study. Further details can be found in the Methods section. **B**, STRING pathway analysis of the 110 genes identified in at least 2 CRISPR screens whose loss sensitizes cells to PARPi treatment. BER: base excision repair; HRR: homologous recombination repair; FA: Fanconi anaemia pathway; RER: ribonucleotide excision repair; DDR: DNA damage response. **C**, Ranking of the top 22 genes identified in our analyses. Genes are colour coded for the different DNA repair pathways they are primarily linked to. Acronyms are as in B. **D**, Top 7 biological processes enriched in the pathway analysis. **E, F**, Dose-response curve for SKOV3 BRCA2 KO (clone 13) (E) and DU145 BRCA1 KO (clone A1) (F) isogenic pairs treated with olaparib for 10-14 days in clonogenic survival assays. Results are shown as mean of n=4 biological replicates ± SD for the dose-response curves.

Our analyses identified *bona fide* clinical biomarkers of sensitivity to PARPi such as *BRCA1* and *BRCA2*. In addition, we also identified others such as the RNASEH2 complex genes (*RNASEH2A, RNASEH2B, RNASEH2C*); *CHD1L*, which encodes the chromatin remodeler ALC1; the gene encoding the nucleotide sanitizer DNPH1 or genes involved in DNA base-excision repair (*XRCC1, POLB, LIG1*), which have already been shown to play roles in PARPi responses that are not dependent on HRR defects (12-14,29-31) (**Fig 1B-C**).

Lack of relevant cellular models has prevented the direct comparison between the *in vitro* responses to PARPi in cell lines deficient in validated clinical biomarkers such as *BRCA1* or *BRCA2* versus other HRR genes. This is particularly the case in cell lines of ovarian and prostate origin. As such, we decided to generate CRISPR-mediated *BRCA1* or *BRCA2* gene knock-outs in the ovarian adenocarcinoma cell line SKOV3 and the prostate carcinoma DU145 cell line (**Supp Fig S1A** and Methods). These cell lines were chosen based on their innate resistance to olaparib treatment, which would make acquisition of sensitivity caused by any given gene KO to be easier to observe, and by them being amenable to colony formation assays, the gold standard to measure PARPi sensitivity *in vitro* (see Methods).

We generated several *BRCA2* KO clones in SKOV3 cells and *BRCA1* KO clones in DU145 cells and validated them functionally to confer sensitivity to olaparib using the colony formation assay (**Fig 1E-F** and **Supp Fig S1B-D**). The IC50 values for olaparib in the *BRCA1* or *BRCA2* KO cell lines we generated (0.067 µM and 0.051 µM, respectively) are in line with previously published data (24), further validating our approach.

### Defects in core HRR factors phenocopy BRCA deficiency

PALB2 and the RAD51 paralogs are key proteins in the HRR pathway and we identified their loss as a cause of sensitivity to PARPi in our CRISPR screen analyses (**Fig 1B; Supp Table S2**). Mammalian cells with mutations in these factors display increased spontaneous chromosomal abnormalities and sensitivity to DNA damaging agents (32,33). Mutations in *PALB2, RAD51B* or *RAD51C* have been explored in clinical trials as biomarkers of sensitivity to PARPi (7,10,11). As such, we generated *PALB2, RAD51B* or *RAD51C* KO clones in the SKOV3 ovarian cell line (**Supp Fig S2A-F**) and compared their sensitivity to olaparib against the *BRCA2* KO SKOV3 cell line. Importantly, *PALB2* or *RAD51C* deficiency phenocopied *BRCA2* loss, while the level of sensitization provided by *RAD51B* inactivation was more modest (**Fig 2A**). We generated an additional *RAD51B* KO in the DU145 prostate cell line (**Supp Fig S2G**) and compared its sensitivity to olaparib against the *BRCA1* KO DU145 cell line. Similar to what we observed in SKOV3, RAD51B inactivation caused a more modest sensitization to olaparib than that observed upon BRCA1 loss (**Fig 2B**). Interestingly, a similar level of sensitization to olaparib in *RAD51B* KO cells was observed upon inactivation in DU145 of the ATPase RAD54L (**Fig 2B, Supp Fig S2H**), whose loss also conferred sensitivity to PARPi in CRISPR screens (**Supp Table S2**).

**Figure 2.**
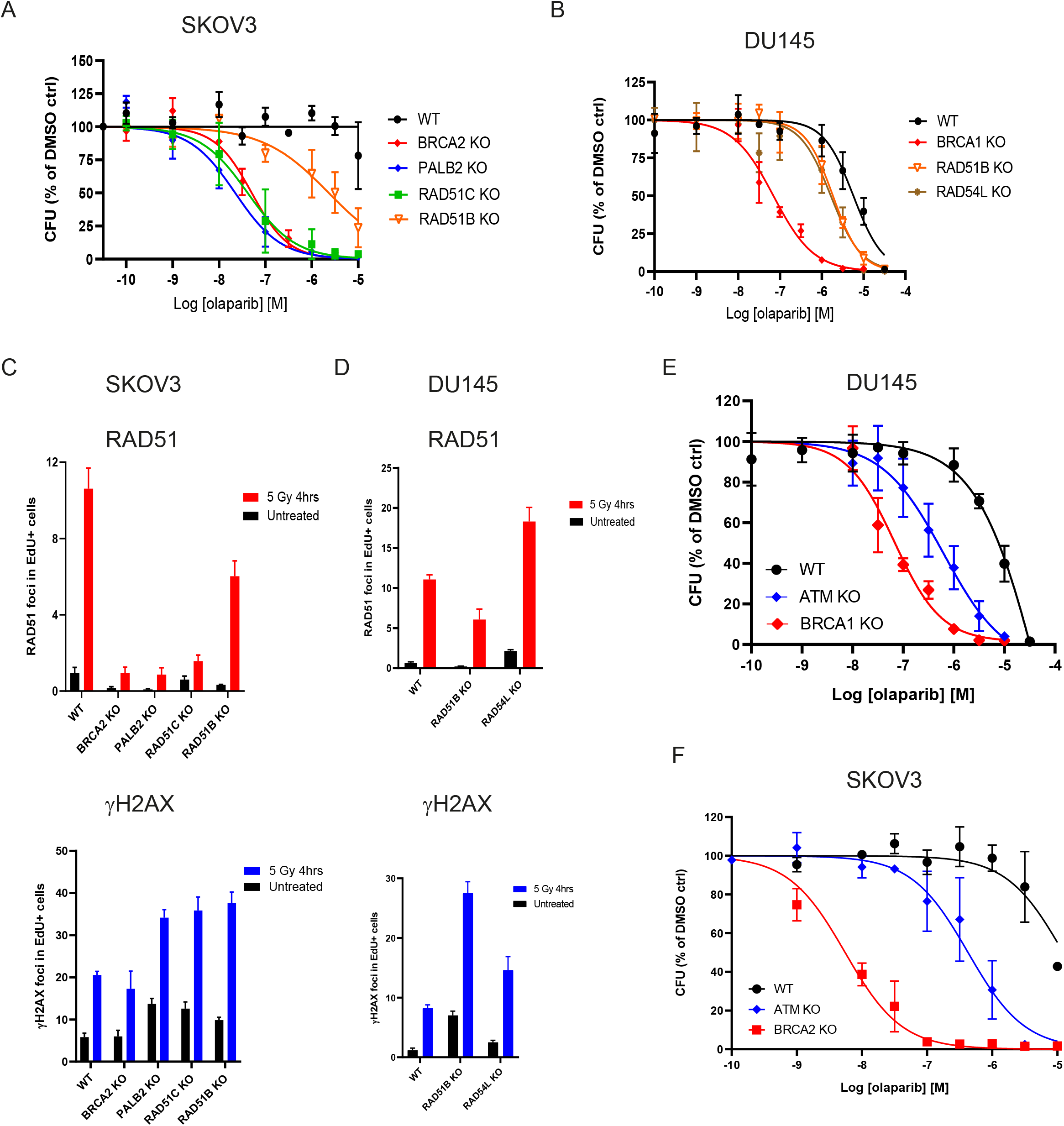
CRISPR/Cas9 knockout of *PALB2, RAD51B, RAD51C, RAD54L* or *ATM* sensitises cell models to olaparib treatment. **A-B**, Dose-response curves for SKOV3 (A) and DU145 (B) isogenic pairs treated with olaparib for 10-14 days in clonogenic survival assays. Results are shown as mean of n=4 biological replicates ± SD. **C-D**, Quantification of RAD51 and γH2AX foci by immunofluorescence in SKOV3 (C) and DU145 (D) isogenic pairs 4 h after treatment with 5 Gy of ionizing radiation or in untreated cells. Results are shown as mean number of foci per nucleus in replicating (EdU-positive) cells (n=3 biological replicates ± SD). **E-F**, Dose-response curves for DU145 (E) and SKOV3 (F) isogenic pairs treated with olaparib for 10-14 days in clonogenic survival assays. Results are shown as mean of n=4 biological replicates ± SD for the dose-response curves. Curves for WT, *BRCA1* KO DU145 and *BRCA2* KO SKOV3 are the same as the ones shown in Figure 1.

To better understand the different responses observed with the several gene KO produced in the canonical HRR pathway, we measured accumulation of the RAD51 recombinase at ionizing radiation-m induced nuclear foci as a surrogate for HRR proficiency (34). Consistent with literature reports, we observed a stark decrease in RAD51 foci formation in the SKOV3 cell lines defective for BRCA2, PALB2 or RAD51C, indicative of inefficient HRR, despite the induction of DNA damage as measured by detection of the phosphorylated form of histone variant H2AX (known as γH2AX; **Fig 2C**).

Importantly, and in agreement with previous reports (32), RAD51B inactivation resulted in a more modest reduction in RAD51 foci formation both in SKOV3 and DU145 cells despite efficient DNA damage induction (**Fig 2C-D**), consistent with the reduced sensitivity of *RAD51B* KO cells to olaparib treatment. Also as previously reported, RAD54L deficiency led to an increase in RAD51 foci formation (**Fig 2D**), which has been linked to the impaired removal of RAD51 molecules upon resolution of HRR and/or the increased number of RAD51 pre-synaptic complexes required to find homology on themdonor DNA strand (35,36).

Collectively, these data provide evidence confirming a similar level of response to olaparib in BRCA2, PALB2 and RAD51C deficient models, a somewhat more modest sensitivity caused by RAD51B or RAD54L deficiencies, and a link between drug response effects to HRR defects in the form of RAD51 foci formation abnormalities.

### *ATM* loss as biomarker for olaparib sensitization

Our analyses of CRISPR screen data identified *ATM* loss as one of the most recurrent events driving sensitivity to PARPi (**Fig 1C** and **Supp Table S2**). Although ATM is not considered a core HRR factor, its deficiency has been linked to sensitivity to PARPi (37,38) and *ATM* mutations have been explored as patient selection biomarkers in clinical trials assessing the efficacy of PARPi (7,10,11). We generated and functionally validated ATM KO cells in DU145 (**Supp Fig S3A-B**) and SKOV3 (**Supp Fig S3C-D**) and compared them with their BRCA mutant counterparts for their response to olaparib. ATM inactivation caused sensitivity to olaparib in both cell lines, however, the sensitization effect did not reach the levels observed for BRCA mutations (**Fig 2E-F**). To confirm these findings, we generated an additional ATM KO in the colorectal carcinoma cell line DLD1 (**Supp Fig S3E**), for which a BRCA2 KO was already available (39). Importantly, DLD1 ATM KO cells were also sensitive to olaparib but less so than their BRCA2 KO counterpart (**Supp Fig S3F**). Given the relatively high prevalence of *ATM* mutations in prostate cancer (7), we decided to generate an additional ATM KO cell line in another prostate cancer cell line, LNCAP (**Supp Fig S3G**), which also resulted in increased sensitivity to olaparib treatment (**Supp Fig S3H**). Taken together, these results confirm that ATM loss confers sensitivity to PARPi in cell lines from a range of different tumour types.

### Clinical prevalence of mutations identifies *XRCC3* loss in prostate cancer

CRISPR LoF screens explore, by definition, responses in a setting where the function of any given gene is completely or almost completely abrogated. As such, we created an analysis pipeline of tumour pan-cancer patient data from The Cancer Genome Atlas (TCGA) to explore the prevalence of biallelic LoF events (including deleterious somatic and germline mutations, homozygous deletions and promoter hypermethylation) in each of the 110 confidence genes in our dataset in the subset of tumour types where PARPi are already available treatment options (ovary, breast, prostate and pancreas; **Fig 3A, Supp Table S3** and Methods). Importantly, our analyses identified biallelic LoF of *BRCA1* or *BRCA2* in 20% and 9% of ovarian cancer samples, respectively, in line with what has been previously reported (**Fig 3B**) (40,41).

**Figure 3.**
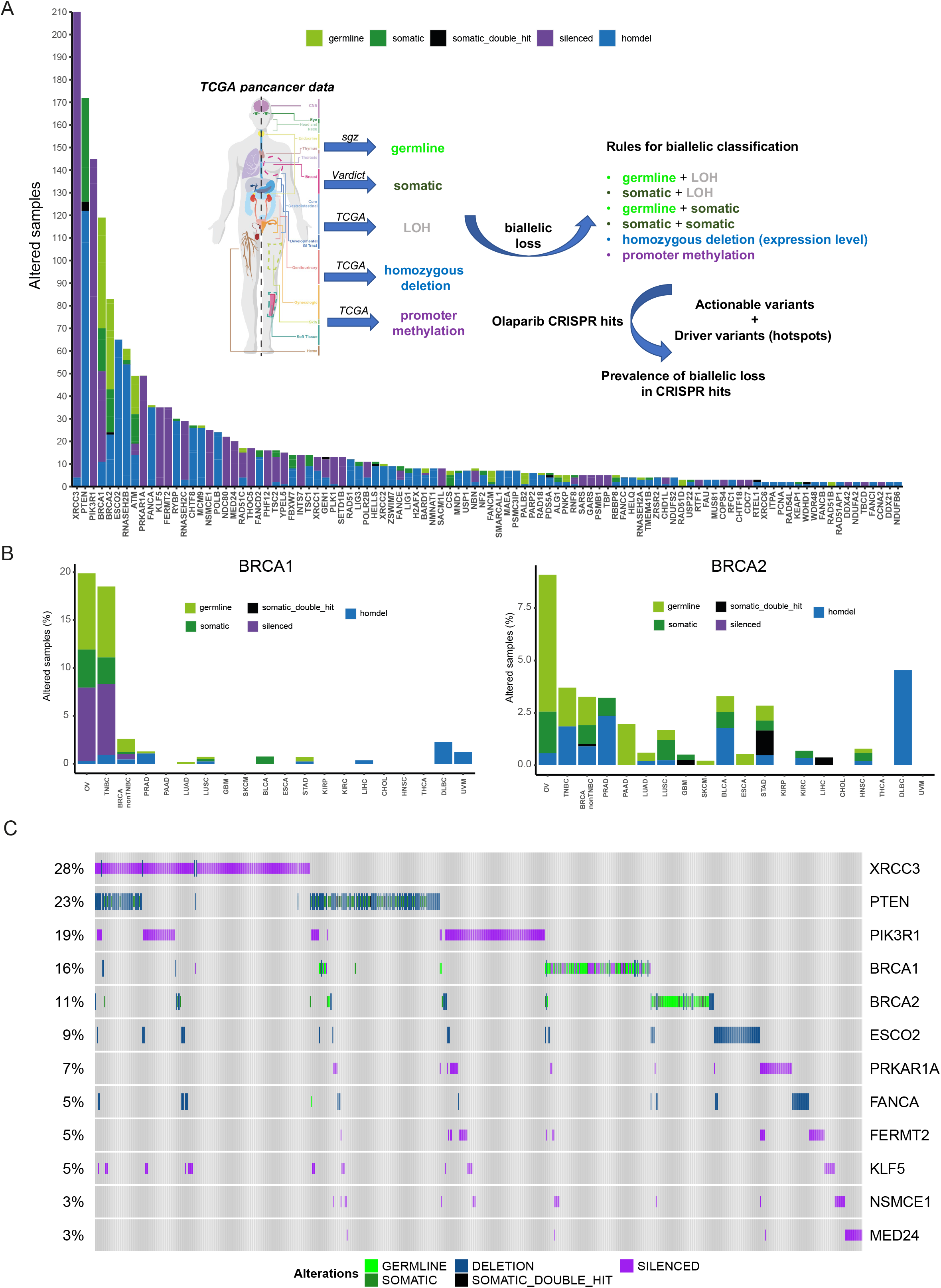
Analyses of genes identified through CRISPR/Cas9 screening in tumour datasets. **A**, Analysis of the total number of samples showing biallelic loss of the 110 confidence genes identified through CRISPR screens in tumour types where PARPi are currently approved (ovarian, breast, pancreas, prostate). The inset shows the pipeline design (see also Methods). **B**, Frequency of biallelic inactivation events for *BRCA1* (left panel) and *BRCA2* (right panel) across the different tumour types analysed. Ovarian (OV), triple-negative breast cancer (TNBC), prostate adenocarcinoma (PRAD), pancreatic adenocarcinoma (PAAD), lung adenocarcinoma (LUAD), lung squamous cell carcinoma (LUSC), glioblastoma (GBM), bladder urothelial carcinoma (BLCA), esophageal carcinoma (ESCA), stomach adenocarcinoma (STAD), kidney renal papillary cell carcinoma (KIRP), kidney renal clear cell carcinoma (KIRC), liver hepatocellular carcinoma (LIHC), cholangiocarcinoma (CHOL), head and neck squamous cell carcinoma (HNSC), thyroid carcinoma (THCA), diffuse large B cell lymphoma (DLBCL), uveolar melanoma (UVM). **C**, Oncoprint of the top 12 genes showing biallelic LoF in the subset of tumour types where PARPi are a treatment option (ovary, breast, pancreas, prostate). Each bar represents an individual tumour. Percentages are for the number of altered samples in the whole dataset.

Interestingly, from the list of 110 confidence genes, *XRCC3* was the most frequently altered gene in our analyses specifically in PARPi-approved settings (**Fig 3A, C**). XRCC3 is one of the five RAD51 paralogs encoded in the human genome (the others being RAD51B, RAD51C, RAD51D and XRCC2) and its function in HRR is well described (32). Accordingly, statistical analyses of the most altered genes in our dataset showed that LoF of *XRCC3* is mutually exclusive with either *BRCA1* (Fisher’s exact p value 0.002) or *BRCA2* (Fisher’s exact p value 0.01) LoF when considering the pan-cancer dataset (**Supp Fig S4A-B**), an effect that becomes even more significant when limiting the analysis to ovarian, breast, pancreatic and prostate tumours (*BRCA1* p value 4.1E-07; *BRCA2* p value 6.4E-04; **Fig 4A**).

**Figure 4.**
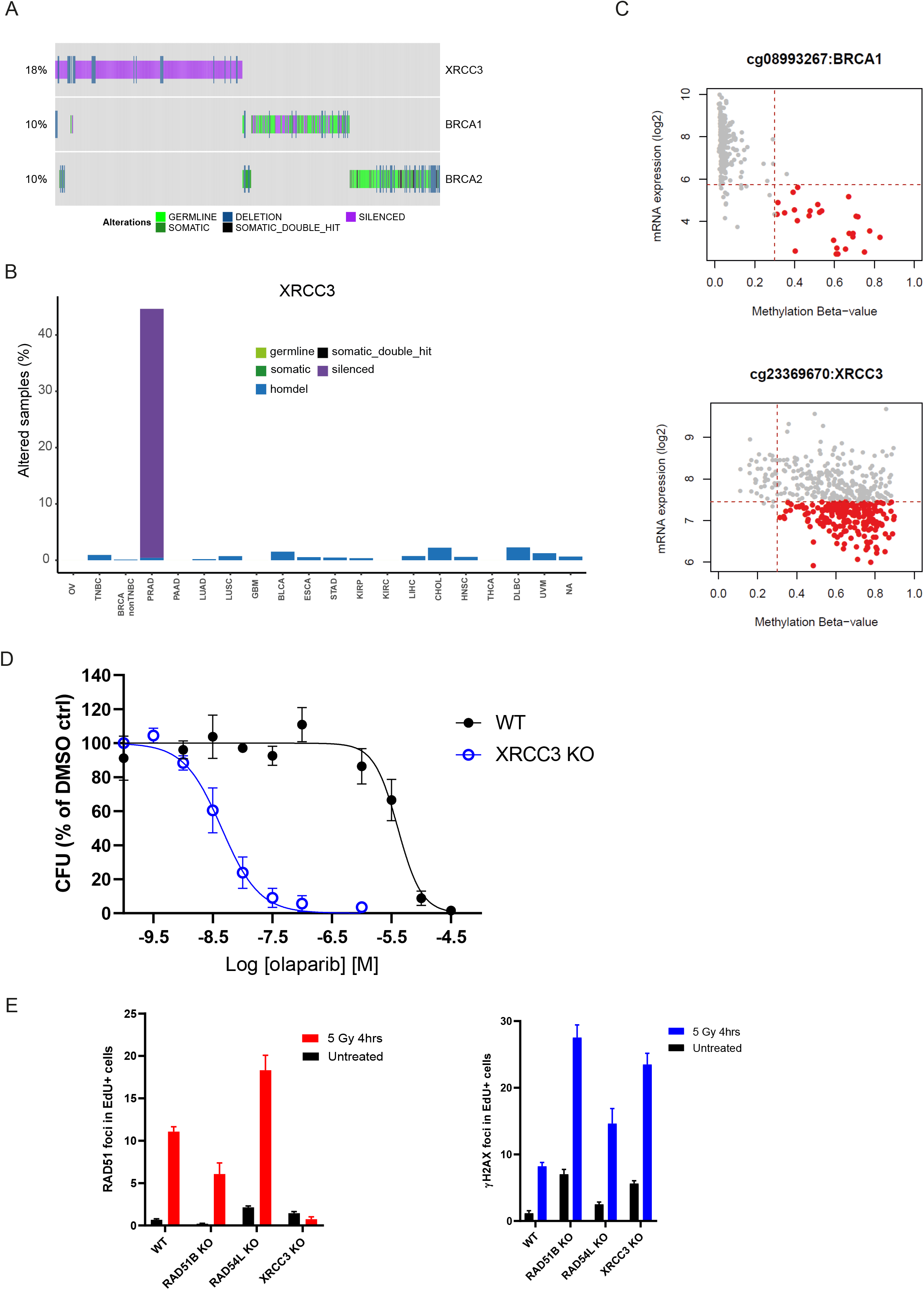
*XRCC3* gene silencing as potential biomarker of olaparib sensitivity in prostate cancer. **A**, Oncoprint depicting mutual exclusivity between LoF of *XRCC3, BRCA1* and *BRCA2* in the pan-cancer analysis. Each bar represents an individual tumour. Percentages indicate the number of altered samples in the whole dataset. **B**, *XRCC3* biallelic inactivation is primarily driven by gene silencing in prostate cancer. Tumour types are the same as listed in Figure 3. **C**, Correlation between *BRCA1* (top panel) and *XRCC3* (bottom panel) mRNA expression and promoter methylation scores. Highlighted in red are samples with high methylation scores and low mRNA expression values. **D**, Dose-response curves for DU145 isogenic *XRCC3* KO cells treated with olaparib for 10-15 days in clonogenic assays. Results are shown as mean of n=3 biological replicates ± SD. **E**, Quantification of RAD51 and γH2AX foci by immunofluorescence in DU145 isogenic pairs 4 h after treatment with 5 Gy of ionizing radiation or in untreated cells. Results are shown as mean number of foci per nucleus in replicating (EdU-positive) cells (n=3 biological replicates ± SD). Data for WT, *RAD51B* and *RAD54L* KO are the same as shown in Figure 2C.

Although there was some level of homozygous deletions adding to the *XRCC3* LoF events in our dataset, it was promoter methylation in prostate cancer samples what drove their prevalence (**Fig 4B-C** and **Supp Table S3**). As a way to mimic this *XRCC3* LoF state, we generated *XRCC3* KO in the DU145 prostate cancer cell line (**Supp Fig S4C**). Strikingly, XRCC3 loss conferred a high level of sensitivity to olaparib in DU145 cells (**Fig 4D**), which correlated with a stark reduction in the ability of these cells to form RAD51 foci upon IR treatment with no impact on γH2AX foci formation (**Fig 4E**). Taken together, our data confirm XRCC3 loss as a bona fide marker of PARPi sensitivity with potential clinical relevance, especially in the prostate cancer setting.

## DISCUSSION

Although a number of functional genomics screens have been performed to explore the genetic determinants of PARPi sensitivity in different cellular models, this study is the first one to try to put the identified biomarkers of sensitivity in a context of prevalence based on available tumour genetic data. We aimed to achieve this goal in two ways. First, by generating head-to-head olaparib sensitivity data on a set of genes identified in the screens, mutations of which have been explored as patient selection biomarkers in PARPi clinical trials (5,7,10,11) and that are involved in different aspects of DNA repair biology, especially HRR. This comparison has been importantly done for the first time in tissue-type relevant cell models (ovary and prostate) with the appropriate clinically-approved benchmark controls for PARPi sensitivity (BRCA1 or BRCA2 deficiency). Together with BRCA1 and BRCA2, deficiency in PALB2 or RAD51C caused the most significant increases in sensitivity to olaparib in these *in vitro* models (**Supp Fig S5**), highlighting the key role these proteins perform in HRR and suggesting that inactivating mutations in *PALB2* or *RAD51C* could be considered as equivalent to *BRCA* mutations with regards to their response to PARPi treatment.

Inactivation of a second group of genes, consisting of *ATM, RAD51B* and *RAD54L*, generated intermediate sensitivity profiles to olaparib treatment when compared to *BRCA1* or *BRCA2* loss in the relevant cell models (**Supp Fig S5**). Our data with *RAD51B* are consistent with published literature (32) and our results with *ATM* are in agreement with the differential responses observed in prostate cancer patients harbouring tumours with mutations in *ATM* treated with olaparib when compared to those carrying *BRCA2* mutations (7). Given that we observed a reduced impact on HRR proficiency (as measured by RAD51 foci formation) in RAD51B or RAD54L deficient cells compared to those lacking BRCA2, PALB2 or RAD51C (**Fig 3C-D**) and that ATM does not play a central role in HRR (38,41), these data suggest that significant impairment of HRR is required to elicit a BRCA-like response to olaparib. Notwithstanding, our data support the inclusion of mutations in these genes as prospective patient selection biomarkers for PARPi treatment, as highlighted in recent prostate cancer clinical trials (7,10,11). For the specific case of ATM, and given the synthetic lethal interaction described between ATM loss and inactivation of the other DNA-damage response apical kinases ATR or DNA-PK (42), it is possible that combination of PARPi with ATR or DNA-PK inhibitors could result in better responses than any single agent approach (43,44).

Our second attempt to provide a more relevant clinical context to the genes identified as potential biomarkers of PARPi response through CRISPR screening was to develop an analysis pipeline assessing the prevalence of biallelic LoF events in a wide-range of tumour types. It was surprising that our analysis identified LoF of *XRCC3* as the most prevalent in PARPi-treated tumour types among the 110 confidence genes, mostly driven by gene silencing caused by promoter methylation in prostate cancer. As expected given the known role of XRCC3 in HRR, its inactivation is mutually exclusive with loss of BRCA1 or BRCA2. In that regard, it is important to highlight that no mutual exclusivity was identified between LoF of bona fide HRR genes such as *BRCA1* or *BRCA2* and genes like *RNASEH2B* or *POLB*. Interestingly, however, a degree of co-occurrence was detected between LoF of the cohesin factor gene *ESCO2* and the BER gene *POLB*, and between *BRCA2* and *RNASEH2B* (**Supp Fig S3B**). It will be interesting to explore whether combinations of these biomarkers could drive better PARPi responses in the relevant tumour types.

XRCC3 forms a functional complex (the CX3 complex) with another RAD51 paralog, RAD51C, whose LoF events are also mainly caused by gene silencing, in this case in breast cancer samples (**Supp Table S3**). RAD51C forms another protein complex (the BCDX2 complex) with other RAD51 paralogs (RAD51B, RAD51D and XRCC2) that does not include XRCC3 (32). Given that XRCC3 deficiency causes a high level of sensitivity to olaparib in the DU145 prostate cancer cell line, and that RAD51C loss phenocopies PARPi sensitivity caused by BRCA2 loss in the ovarian cell line SKOV3, while the same is not true for the other BCDX2 complex component RAD51B, it is tempting to speculate that there is a more important function of the CX3 RAD51 paralog complex in mediating PARPi responses. Whatever the case, the identification of *XRCC3* silencing as an important LoF event in prostate cancer opens the possibility to explore its clinical relevance in a tumour type where olaparib is already a treatment option. Retrospective analysis of tumour material linked to patient outcome in ongoing prostate cancer clinical trials (45,46), once available, could definitely shed light into the importance of this newly identified biomarker.

## Authors’ Disclosures

KJ, AG, JA, GI, SB, JH, DB, MA, MJO, EL and JVF are or were employees of AstraZeneca at the time of conducting these studies. Several authors hold stock or shares in AstraZeneca.

## Acknowledgements

This study was funded by AstraZeneca. We thank Dr David Fisher and his team for providing some of the KO models used in this study.

## Authors’ contributions

JVF, EL and MJO conceived the study and designed the research plan with KJ, AG, GI and JA. KJ and AG performed all other in vitro work with technical support from GI, SB and JH. JA performed the bioinformatics analysis from TCGA data with advice from JVF. Analyses of CRISPR datasets were performed by DB and MA. All authors contributed to data interpretation. KJ, AG, JA and JVF prepared the figures and tables, and KJ and JVF wrote the manuscript. All authors reviewed and approved the final manuscript.

## Functional Genomics Centre members

Douglas Ross-Thriepland, David Walter, Khalid Saeed, Jennifer Hillis, Adam Spruce, Mel Lad, Abigail Shurr, Dave Kim, Curtis Hart, Rebecca England, Jennifer Muscat, Sebastian Lukasiak, Angelos Papadopoulos, Alex Kalinka, Marica Gaspari, Daniel Barrell, Miika Ahdesmäki, Lu Li, Maryam Ghaderi, Carolina Florez Zapata, Ultan McDermott, Greg Hannon.

## Notes

**Conflict of interest statement:** KJ, AG, JA, GI, SB, JH, DB, MA, MJO, EL and JVF are or were employees of AstraZeneca at the time of conducting these studies. Several authors hold stock or shares in AstraZeneca.

