## Supplementary Figures and legends for "Drug-gene interaction screens coupled to tumour data analyses identify the most clinically-relevant cancer vulnerabilities driving sensitivity to PARP inhibition"

### Supplementary Figure 1

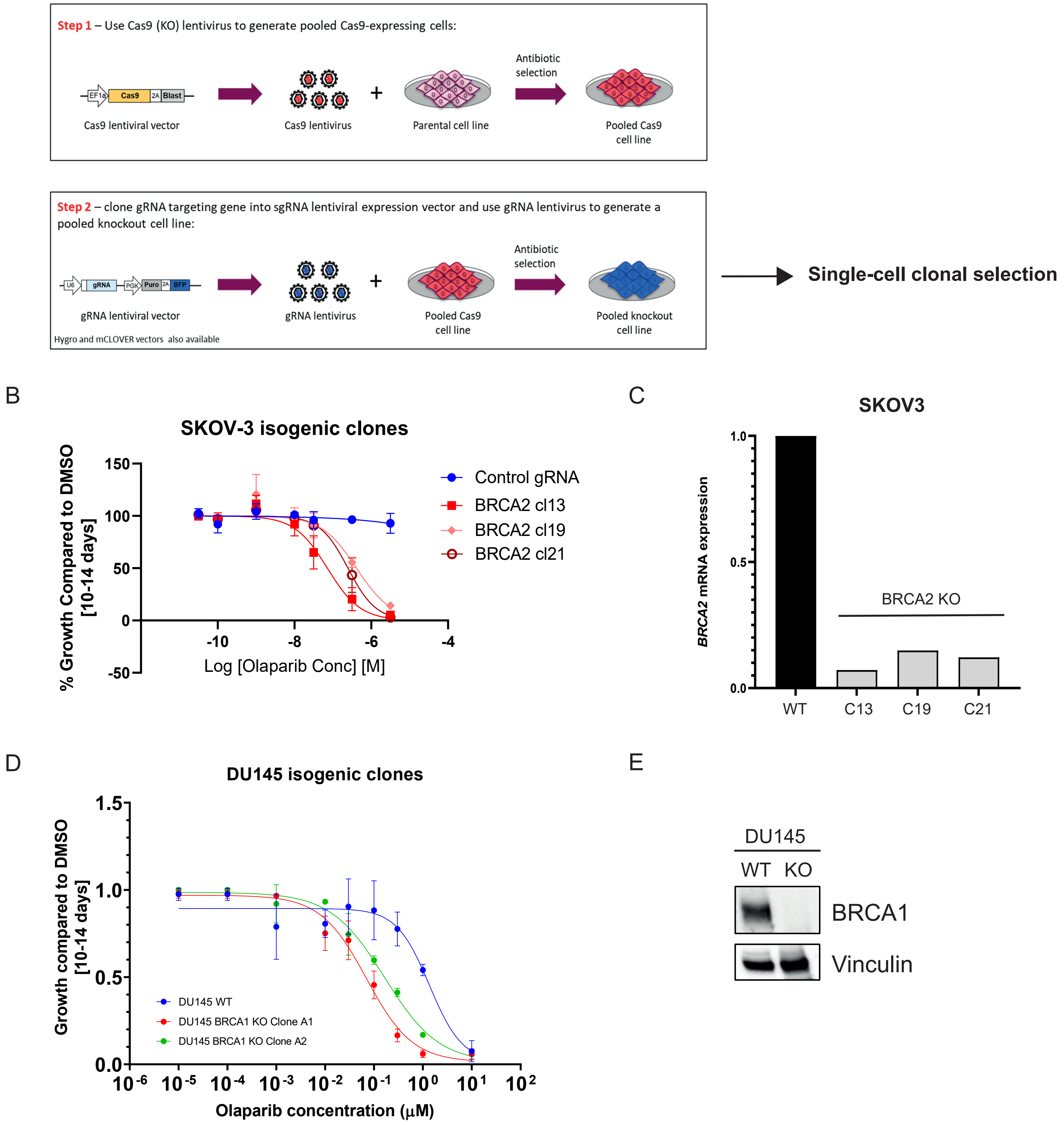

**Supplementary Figure S1. A,** Workflow to generate isogenic HRR KO cell models (see also Methods). **B,** Dose-response curve for SKOV3 BRCA2 KO isogenic pairs treated with olaparib for 10-14 days in clonogenic survival assays. Results are shown as mean of n=4 biological replicates ± SD for the dose-response curves. Clone 13 was selected for further experimentation. **C,** Fold-change mRNA expression in WT and KO cells to assess BRCA2 loss in SKOV3 cells. **D,** Dose-response curve for DU145 BRCA1 KO isogenic pairs treated with olaparib for 10-14 days in clonogenic survival assays. Results are shown as mean of n=2 biological replicates ± SD for the dose-response curves. Clone A1 was selected for further experimentation. **E,** Western blot data to assess BRCA1 loss in DU145 cells (clone A1).

### Supplementary Figure 2

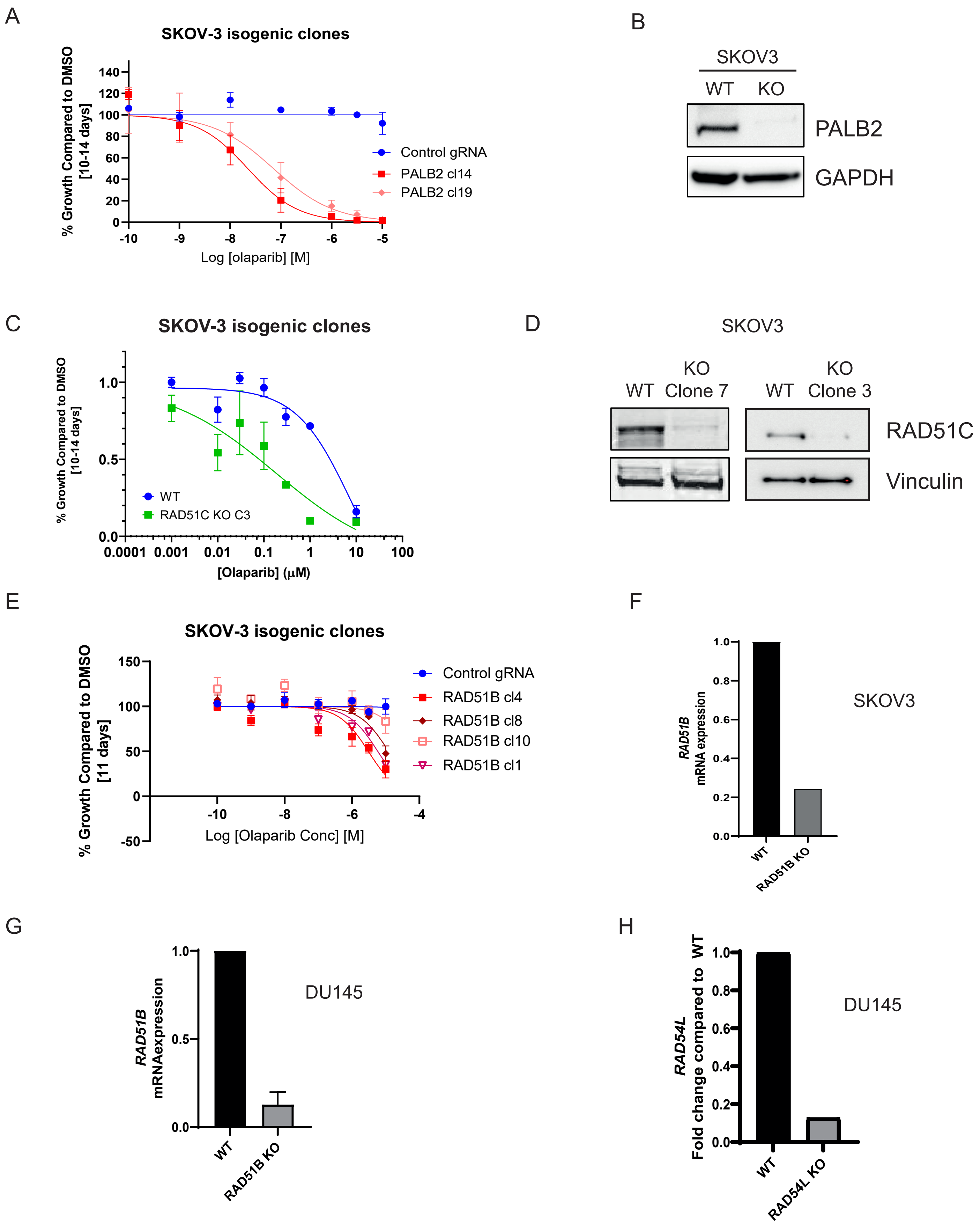

**Supplementary Figure S2. A,** Dose-response curve for SKOV3 PALB2 KO isogenic pairs treated with olaparib for 10-14 days in clonogenic survival assays. Results are shown as mean of n=4 biological replicates ± SD for the dose-response curves. Clone 19 was selected for further experimentation. **B,** Western blot data to assess PALB2 loss in SKOV3 cells (clone 19). **C,** Dose-response curve for SKOV3 RAD51C KO clone 3 isogenic pairs treated with olaparib for 10-14 days in clonogenic survival assays. Results are shown as mean of n=3 biological replicates ± SD for the dose-response curves. **D,** Western blot data to assess RAD51C loss in SKOV3 cells (clone C7 was selected for further experimentation and its dose-response curve is shown in Figure 2A). **E,** Dose-response curve for SKOV3 RAD51B KO isogenic pairs treated with olaparib for 10-14 days in clonogenic survival assays. Results are shown as mean of n=4 biological replicates ± SD for the dose-response curves. Clone 4 was selected for further experimentation. **F-H,** Fold-change mRNA expression in WT and KO cells to assess RAD51B loss in SKOV3 cells (clone 4) (F) and DU145 cells (G), and loss of RAD54L in DU145 cells (H).

### Supplementary Figure 3

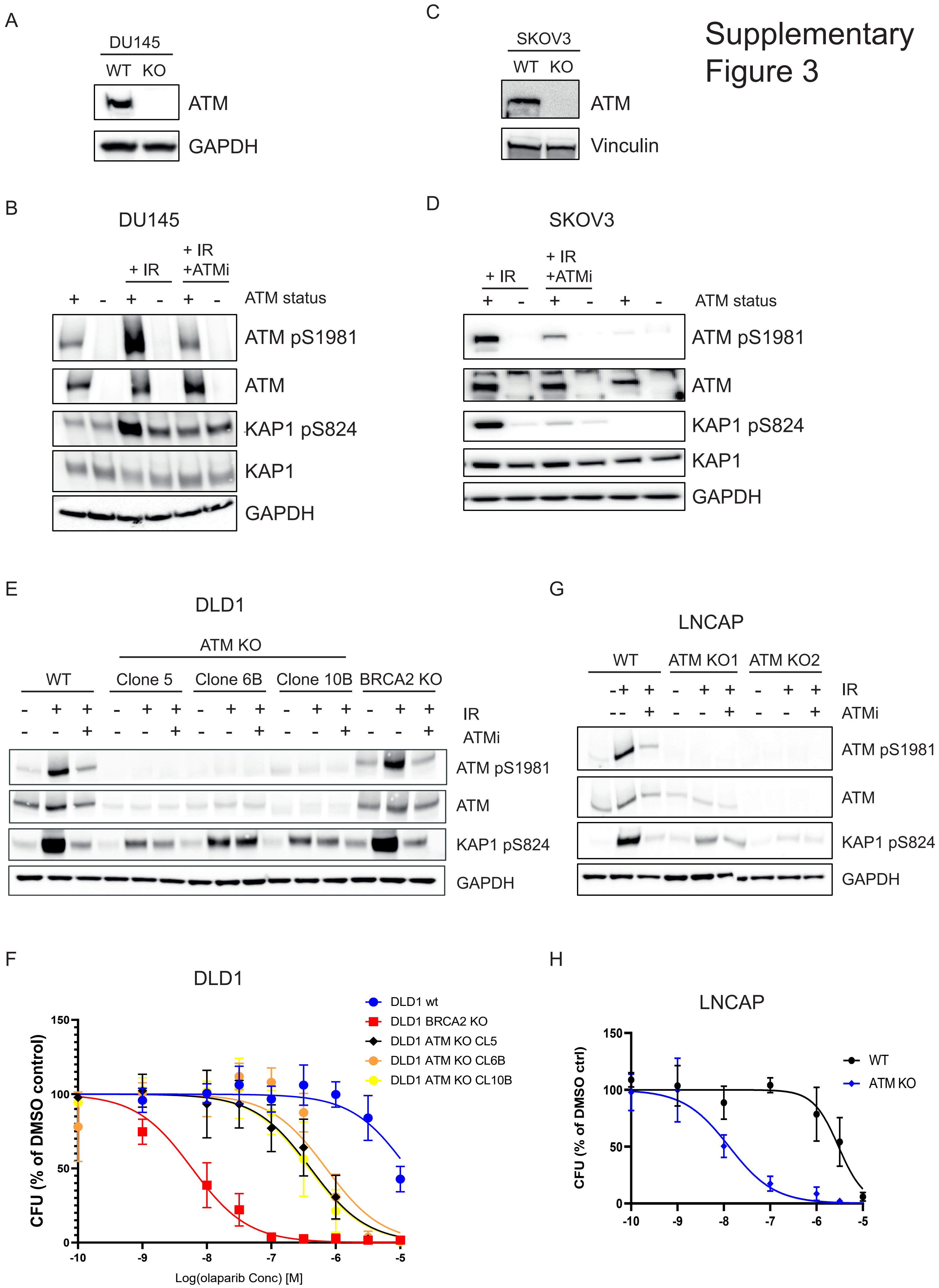

**Supplementary Figure S3. A-E and G,** Western blots to assess ATM loss (A, C) and ATM signalling (B, D, E, G) 4 h after treating cells with 5 Gy of ionizing radiation (IR) in the presence or absence of ATM inhibitor (AZD0156, 100 nM) added 2 h before treatment in DU145 (B), SKOV3 (D), DLD1 (E) and LNCAP (G) *ATM* KO isogenic cell lines. **F and H**, Dose-response curves for ATM KO DLD1 clones (F) and LNCAP (H) isogenic pairs treated with olaparib for 10-14 days in clonogenic survival assays. Results are shown as mean of n=4 biological replicates ± SD for the dose-response curves. Clonogenic survival data on (H) was generated with LNCAP ATM KO2 cells (see panel G). Clone 5 of the DLD1 ATM KO cells was selected for further experimentation.

### Supplementary Figure 4

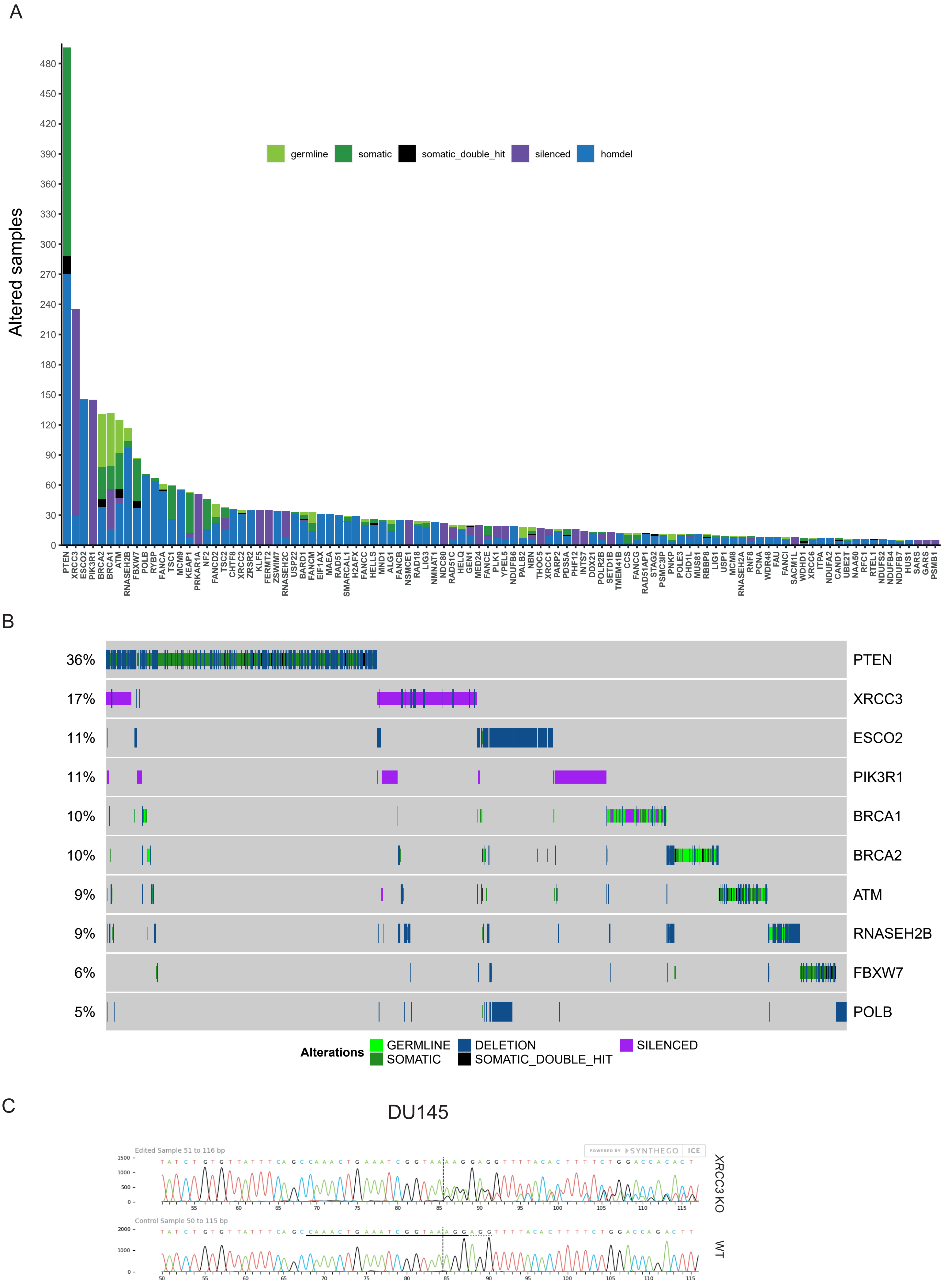

**Supplementary Figure S4. A,** Pan-cancer analysis of the total number of samples showing biallelic loss of the 110 confidence genes identified through CRISPR screens. **B,** Oncoprint of the top 10 genes showing biallelic LoF in the pan-cancer analysis. Each bar represents an individual tumour. Percentages are for the number of altered samples in the whole dataset. **C,** Inference of CRISPR Edits (ICE) analyses of the edited region of the *XRCC3* gene in WT (bottom) or *XRCC3* KO (top) DU145 cells. ICE quantification reported an overall editing efficiency of 98% in the KO cells, with a 1 bp deletion accounting for 22% of the events and a 1 bp insertion accounting for the remaining 77%.

### Supplementary Figure 5

| | Cell line olaparib IC50 (μM) | | |
|---|---|---|---|
| **Genotype** | SKOV3 | DU145 | DLD1 |
| Parental | > 10 | 8 | > 10 |
| ATM KO | 1 | 0.45 | 0.381 |
| BRCA1 KO | | 0.067 | |
| BRCA2 KO | 0.051 | | 0.006 |
| PALB2 KO | 0.0223 | | |
| RAD51B KO | 3 | 1.8 | |
| RAD51C KO | 0.04 | | |
| RAD54L KO | | 1.6 | |
| XRCC3 KO | | 0.007 | |

**Supplementary Figure S5.** Half-maximal inhibitory concentration (IC50) for olaparib in the different isogenic cell lines used in this study as measured in clonogenic survival assays. Values were calculated from the dose-response curves in Figures 1, 2 and 4 and Supplementary Figure S3.
